# Multiplexed dopamine neurons predominate in the ventral midbrain of young macaques

**DOI:** 10.64898/2026.06.03.729859

**Authors:** E.A. Kelly, I. Mahoui, J.L. Fudge

**Author notes:** Corresponding author Del Monte Institute for Neuroscience, University of Rochester Medical Center, 601 Elmwood Avenue, Rochester, NY 14642-0001.

## Abstract

Dopamine (DA) is important in many fundamental behaviors, including positive and negative reinforcement, incentive salience, and decision-making. This behavioral diversity is now known to be due, in part, to neurotransmitter diversity, based on rodent models. To address DA neuron transmitter properties in higher species, we examined the ventral midbrain in 5 young macaques (3 male, 2 females, 3-6 years) using RNAscope *in situ hybridization* for tyrosine hydroxylase (TH), vesicular glutamate transporter 2 (VGluT2) and glutamic acid dehydroxylase 1 (GAD1) across the A10 (VTA; ‘midline VTA’ nuclei and parabrachial nucleus; PBP), the A9 (substantia nigra, pars compact, SNc) and the A8 (retrorubral field, RRF) subregions. We followed up with immunocytochemical studies in the same cohort to infer extent of mRNA and protein matches. There were 7 mRNA phenotypes, with TH-mRNA containing cells forming the largest proportions of all neurons, as expected. Surprisingly, multiplexed TH+ neurons were much more frequent than TH-single labeled neurons overall (TH-VGluT2, 22% and TH-VGluT2-GAD1, 23% compared to TH-single labeled neurons, 19%). GAD1 mRNA co-expression mainly occurred in ‘triple labeled’ cells, i.e. those with VGluT2- and TH mRNA expression. VGluT2 mRNA single-labeled neurons represented only 8%, and GAD1 mRNA single-labeled neurons comprised 20%, of the total population. Proportions of cellular phenotypes were similar across the A10-A9-A8 subregions. Most DA neurons in the young macaque contain multiple transmitters, indicating an important role for fast synaptic transmission alongside dopaminergic transmission in all subregions. We discuss the developmental and circuit implications of these findings in higher primates.

## Introduction

The ventral midbrain houses the most prominent population of dopamine (DA; tyrosine hydroxylase [TH] expressing) cells in the brain ^1, 2^, and composed of three anatomical groups: the ventral tegmental area (VTA, A10), including the midline nuclei and parabrachial pigmented nucleus (PBP, A10), the substantia nigra pars compacta (SNc, A9) and the retrorubral field (RRF, A8)^3-8^. The neurotransmitter DA is a fundamental signaling factor in numerous behaviors including positive and negative reinforcement ^9-11^, decision making ^12^, working memory ^13-15^, incentive and stimulant-induced salience ^16-18^ and purposeful movement ^19^. A diversity of DA and non-DA cell types in the ventral midbrain is now proposed to mediate specific circuitry supporting these different functions ^6, 20, 21, 22^ .

DA neurons are now recognized to release DA with either glutamate ^23, 24^ or GABA ^25^ in some circuits, and glutamate neurons that contain both glutamate and GABA are also recognized in rodent models ^26^. Unsurprisingly, it is these ‘multiplexed’ populations of neurons have garnered immense interest due to the enhanced computational capacity that may arise. Indeed, the heightened complexity regarding the activation of multiple postsynaptic receptor types is balanced by the multi-modal effects of pre- and postsynaptic neuronal modulation, variations in co-release and co-transmission, and differential release in time and space; all parameters that will change the functional outcome ^27^.

There is little information on the extent or distribution of ‘multiplexed’ DA populations in higher species^28, 29^. Since many neuropsychiatric and degenerative illnesses in the human involve alterations in the midbrain DA system, a model that better approximates the human DA system is needed. While the main DA subregions are generally conserved across species ^30^, variations in relative size of the subregions, the interdigitation of the A9 neurons deep into the GABAergic substantia nigra reticulata (SNr), and a large mediolateral extension of the A10 and A8, with fewer clear boundaries across regions are features of human and nonhuman primate^5, 31, 32^.

To update existing models of DA neurotransmitter co-localization in nonhuman primates, we investigated the mRNA markers of transmitters in ventral midbrain neurons in young male and female macaques. To overcome past issues with mRNA detection, we used RNAscope techniques to detect TH, vesicular glutamate transporter-2 (VGluT2), and glutamate decarboxylase-1 (GAD1) mRNA across five animals, and found 7 neuronal phenotypes. Surprisingly, multiplexed TH-containing neurons were the rule rather than the exception for the entire ventral midbrain, a feature that was found across all the DA subregions. In addition, a subpopulation of TH+VGluT2+ labeled neurons also co-contained GAD1 message (TH+VGluT2+GAD1+ phenotype) suggesting a potential mechanism for converting glutamate to GABA. Results are discussed with respect to potential developmental and circuit relevance of large numbers of ‘multiplexed’ neurons in young animals.

## Materials and Methods

### Design

The ventral midbrain comprising the A10, A9, and A8 regions was defined based on morphological and immunohistochemical staining of TH neurons through their rostrocaudal extent. The classic DA subgroup boundaries in monkey and human are drawn according to both TH immunoreactivity (IR) and for expression of the calcium binding protein (CaBP; a marker of DA neurons in the A10 and A8 regions, absent in the A9 region) ^33-36^. The A10 region was further subdivided into the midline VTA nuclei (mVTA) and the large parabrachial pigmented nucleus (PBP). The midline VTA nuclei are composed of the rostral linear nucleus (RLi), the caudolinear nucleus (CLi), the intrafascicular nucleus, the ventral tegmental nucleus (VTA) and paranigral nucleus ^32, 37^. The PBP is dorsal and lateral to the midline VTA nuclei and extends across the entire mediolateral expanse of the midbrain, over the A9 region, to merge with the A8 neurons caudally ^36^.

### Animals

We used brains from three young male and 2 young female monkeys (*Macaque fascicularis*) that had been perfused in 4% paraformaldehyde (PFA). To conserve animals, these cases had received tracer injections in other brain regions as parts of other studies (**Table 1**). We selected cases of young macaques, 3-4 years of age (equivalent to 15-16 human years and 1 seven-year-old male (equivalent to 21-28 human years) ^38^. We previously conducted baseline cortisol measures in a subgroup of these animals, and found little variation among levels or animals over a six-month period, suggesting a stable living environment^39^. The entire brain was coronally sectioned on a freezing sliding microtome at 40 μm and stored in serial compartments in cryoprotectant solution (30% sucrose and 30% ethylene glycol in phosphate buffer (PB)) at -20^°^C.

**Table 1.**
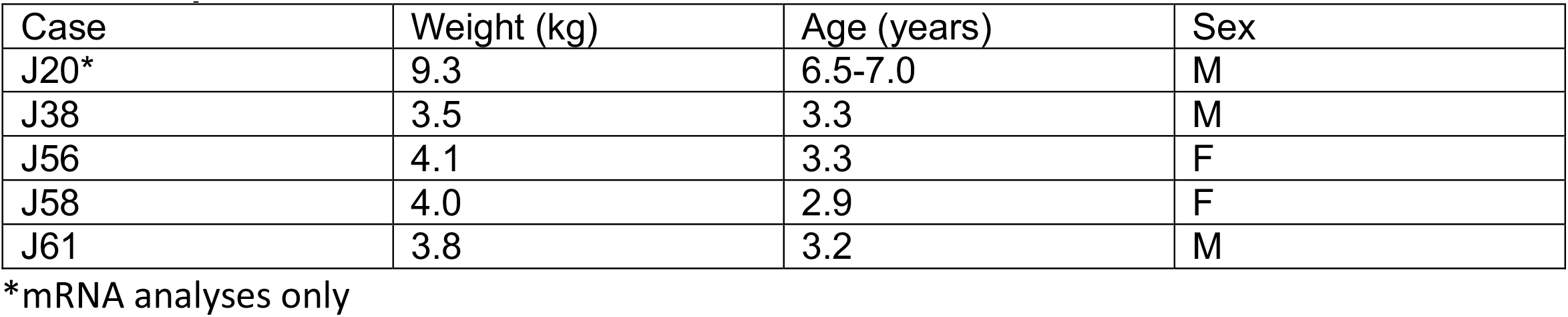
Experimental Cases.

### RNAscope Processing

1:24 hemi-sections through the midbrain from the 3 males (J20, J38, J61) and 2 females (J56, J58) were assessed for TH, VGluT2, and GAD1 transcripts in the DA subregions. Target probes were designed for a 3-plex fluorescent assay for macaque (Mmu) as follows. *Mmu-TH-C1* (tyrosine hydroxylase transcript variant X1 mRNA), *Mmu-SCL17A6-C2* (VGluT2, Macaca mulatta solute carrier family 17, sodium dependent inorganic phosphate cotransporter, member 6 transcript variant 2 mRNA) and *Mmu-GAD1-C3* (Macaca mulatta glutamate decarboxylase 1 mRNA) (Advanced Cell Diagnostics, ACD Bio, Newark CA). Sections were thoroughly rinsed in 0.01M phosphate buffered saline (PBS), then the midbrain was blocked, excised, and mounted onto Superfrost plus slides and stored at -80^°^C until ready for processing. Multiplex RNAscope experiments were processed using RNAse-free reagents on-slide using RNAscope detections kits and reagents following the manufacturer’s protocol with slight modification to improve adhesion of thicker tissue ^40^. Sections were allowed to thaw and air dry, then washed for 10 minutes in distilled, deionized water (ddH_2_O), then postfixed at 4^°^C in 4% PFA, rinsed in 0.01M PBS and gradually dehydrated in ascending concentrations of ethanol. Sections were then baked in a HybEZ oven to reinforce adhesion, treated with 1% H_2_O_2_, rinsed and baked again. Remaining steps were as described in the ACDBio RNAscope protocol (Document # 323100-USM). Following pretreatment, slides were air dried and incubated in protease III followed by a 1:50 dilution of target probes (1-part C2/C3: 50 parts C1) for 2 hours at 40^°^C. Slides were then processed using the RNAscope Multiplex Fluorescent Assay kit version 2 (ACD Bio). The following Opal Dyes were used: Opal 520 (FP1487001KT; Akoya Biosciences), Opal 570 (FP1488001KT) and Opal 690 (FP1497001KT). Sections were washed in Trueblack (1:20K; Biotium, Cat # 23007) in 70% ETOH for 45 sec at room temperature, then coverslipped using Prolong Gold (Invitrogen, Waltham, MA).

### Immunocytochemistry (ICC)

*TH and GAD1*. To investigate the extent to which RNA transcript expression translates into protein expression, near adjacent compartments of tissue for cases 38, 56, 58, and 61 were selected for double labeling for antibodies to TH (1:10K; mouse, MAB318, Millipore, clone LNC1) and GAD1 proteins (1:10K, goat, AF2086, R&D Systems). Anti-tyrosine hydroxylase (TH, mouse, MAB318, Millipore, clone LNC1) specificity is well-documented ^41^, and our results are in agreement with previously reported distribution and cytoarchitecture of TH-immunoreactivity (IR) cells in monkey and human ^3^. Anti-glutamic acid decarboxylase 1 (GAD1/67, goat, AF2086, R&D Systems) targets the enzyme that catalyzes the conversion of glutamate into GABA, is expressed constitutively, and has been characterized in the nonhuman and human primate midbrain ^37, 42, 43^.

Prior to immunostaining, hemi-sections were photobleached to eliminate autofluorescence due to lipofuscin ^44^. After overnight overnight rinses in 0.1M phosphate buffer with 0.3% Triton-X (PB-TX), free-floating sections are submerged in PB in a Petri dish under a direct 6000 luxe light (Husky HD12000DIM) for 48 hours at 4^°^C. Photobleaching eliminated all endogenous fluorescent signal. Hemi-sections from 1:24 compartments were then processed using free-floating methods, blocking tissue in 10% normal donkey serum (NDS) in PB-TX followed by incubation in pooled primary antibodies (TH, 1:10K, mouse, MAB318, Millipore, clone LNC; GAD1, 1:10K, goat, AF2086, R&D Systems) for 96 hours at 4^°^C. Tissue was then rinsed with PB-TX for 6 x 15 minutes, blocked with 10% NDS-PB-TX and incubated with pooled secondary antibodies: donkey anti-mouse 488 (for TH, 1:200, AlexaFluor 488, Invitrogen) and donkey anti-goat 647 (for GAD1, 1:200, AlexaFluor 647, Invitrogen) overnight at room temperature. Following thorough rinsing in PB, tissue was mounted out of 0.1M PB, pH 7.2, and cover-slipped with Prolong Gold anti-fade mounting media (Invitrogen).

#### Calbindin-D28k (CaBP)

Near-adjacent sections to RNAscope and ICC-labeled sections were chosen for permanent CaBP immunostaining to ascertain DA subregions. Tissue compartments were thoroughly rinsed overnight in PB-TX, blocked in 10% NGS, and incubated in CaBP anti-serum (1:10,000, C9848, made in mouse, Sigma, St. Louis, MO) for 96-hours. On day 4, after thorough rinsing and blocking, sections were incubated in a biotinylated secondary antibody (goat-anti mouse, 1:200, Invitrogen) for 40 minutes, followed by more rinses in PB-TX. Immunoreactivity was visualized using an avidin-biotin reaction (Vector Elite kit) and colorized with 0.03 % H_2_O_2_2% and 3’3-diaminobenzidine. Sections then were rinsed, mounted onto gelatin-coated slides, dried, and coverslipped with Permount (Fisher Scientific, Waltham, MA).

### Image Acquisition and Analysis

#### Confocal image acquisition

RNAscope and dual immunofluorescent processed hemisections were captured in sequential channels on a Nikon A1RHD Laser Scanning Confocal microscope using NIS-Elements software (Center for Advanced Microscopy and Nanoscopy). The following excitation (ex) lasers and emission (em) were used for each channel: RNAscope: GAD1-Opal 520, ex 488, em 525/50; TH-Opal 570, ex 561, em 595/50; VGluT2-Opal 690, ex 640, em 650LP. ICC: TH-AlexaFluor 488, ex 488, em 525/50; and GAD1-AlexaFluor 647, ex 640, em 650LP. After optimization, the illumination parameters for each channel were held constant for all sections. A tiled ROI of the ventral midbrain was then collected using a 10X/0.4 NA Plan Apochromat objective.

#### Light microscopic mapping of CaBP-IR

CaBP-IR images were collected on a BX51(Olympus) bright field microscope using a 2X PlanApo objective (2X/0.08NA) with SpotCam acquisition software, then imported into Adobe Illustrator files. The boundaries of CaBP-positive neurons (demarcating mVTA/A10, PBP/A10 and RRF/A8) and CaBP-negative neurons (demarcating SNc/A9) were manually traced using Neurolucida software (Microbrightfield Bioscience, Williston, VT), along with landmarks such as major fiber tracts and blood vessels.

#### RNAscope and ICC Analysis

Immunofluorescent transcript and protein labeling counts were analyzed in conjunction with CaBP boundary overlays using Neurolucida software. Individual micrographs were imported, and manual CaBP boundary overlays for each DA subregion were carefully overlaid, aligning fiducial markers. Each DA subregion on the section was collected separately. Prior to final data collection, iterative trials of cell detection parameters were done in all subregions until there was 100% agreement between automatically selected cells, and validation during random spot checks under high magnification.

After optimization of each channel individually, semi-automated neuronal counts were performed using the “Detect Cells” pipeline. Neuronal inclusion was based on the following “cell strength” settings: *cells size range*; determined by manually measuring the diameter of the ‘largest’ and ‘smallest’ cells, *cell strength and filter range*; manually adjusted inclusion thresholding parameters to only include cells and minimize background artifact. Markers were selected for each primary transcript/protein(e.g. a separate marker for TH, GAD1 and VGluT2 in RNAscope sections) and placed automatically in separate channels. After marker placement, the ‘colocalization’ tool was used to create a new marker for distinct marker combinations. For this step, the ‘marker overlap’ distance was set to the larger cell size setting determined during Neuronal Inclusion steps. Finally, marker placement and colocalization accuracy within each image (and channel) was randomly spot-checked under higher magnification. Individual marker counts were recorded for each neuronal combination and analyzed for each DA subregion across 4-6 sections and summed. The substantia nigra pars reticulata rostromedial tegmental nucleus were excluded from the analysis.

#### Comparing mRNA and protein expression

We compared mRNA and protein expression in animals 38, 56,58, and 61. Total counts for each cellular phenotype were examined to determine whether the relative total expression of TH and GAD within each were similar (**Tables 2 and 3**). While glutamatergic neurons could not be counted in the ICC sections, for comparison across the two studies we later recalculated proportions by subtracting out the VGluT2-only phenotype from the denominator in the mRNA studies.

**Table 2.**
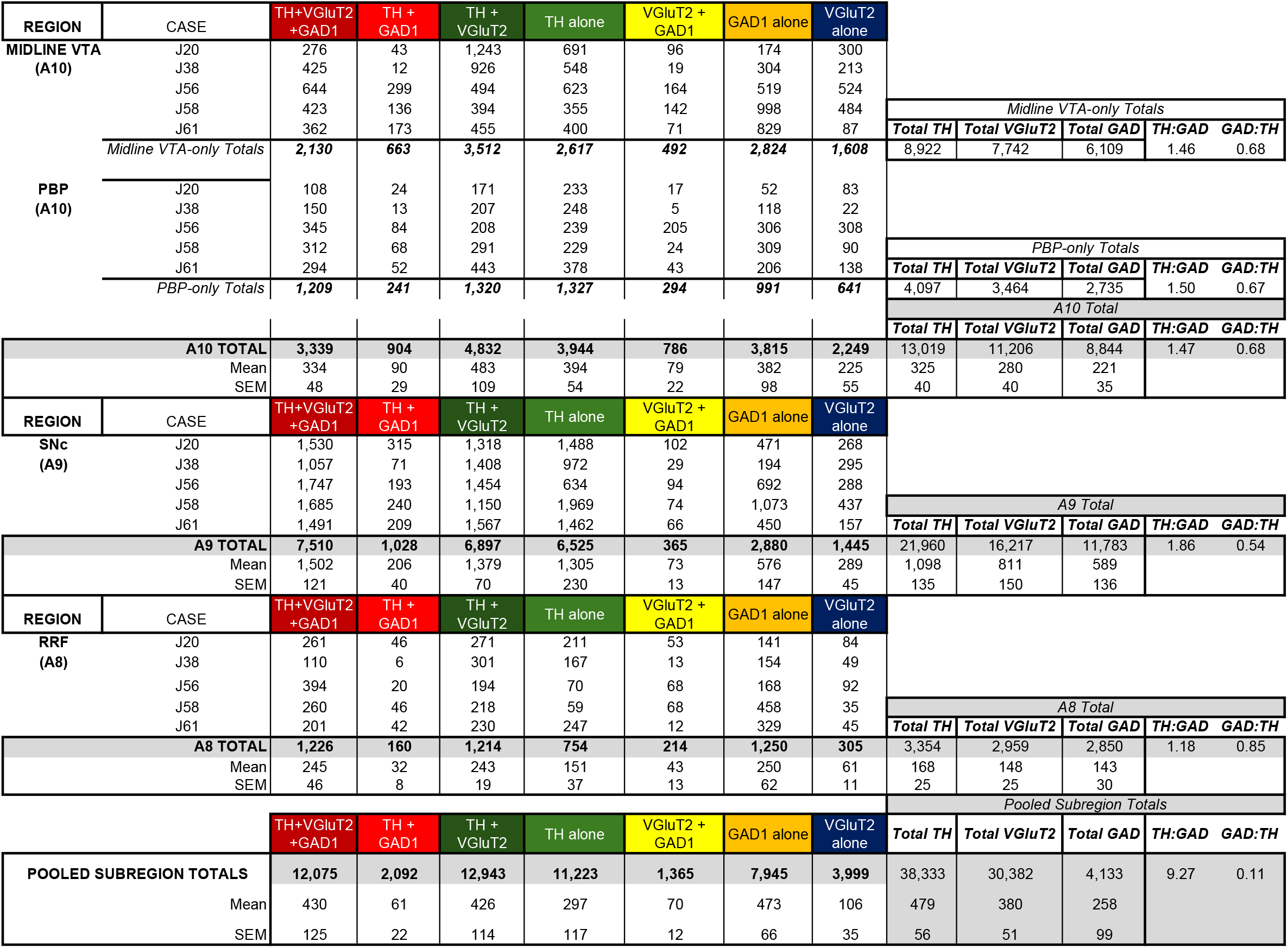
Raw data counts for each subregion in all cases. The midline VTA and PBP comprise the A10 subregion, the SNc is the A9 subregion, and the RRF is the A8 subregion.

**Table 3.** Raw data counts for each subregion in all cases. The midline VTA and PBP comprise the A10 subregion, the SNc is the A9 subregion, and the RRF is the A8 subregion.

| REGION | CASE | Total TH | Total GAD1 | TH+GAD1 | TH alone | GAD1alone |  |  |
| --- | --- | --- | --- | --- | --- | --- | --- | --- |
| MIDLINE VTA<br>(A10) | J38 | 801 | 434 | 315 | 486 | 119 |  |  |
|  | J56 | 581 | 430 | 215 | 366 | 210 |  |  |
|  | J59 | 624 | 365 | 241 | 383 | 124 |  |  |
|  | J61 | 875 | 467 | 319 | 556 | 148 |  |  |
|  | *Midline VTA-only Totals* | 2881 | 1696 | 1090 | 1791 | 601 | *TH:GAD* | *GAD:TH* |
|  |  |  |  |  |  |  | 1.70 | 0.59 |
|  |  | Total TH | Total GAD1 | TH+GAD1 | TH alone | GAD1alone |  |  |
| PBP<br>(A10) | J38 | 438 | 243 | 197 | 241 | 46 |  |  |
|  | J56 | 494 | 308 | 213 | 281 | 95 |  |  |
|  | J59 | 474 | 358 | 256 | 218 | 102 |  |  |
|  | J61 | 636 | 347 | 301 | 335 | 46 |  |  |
|  | *PBP-only Totals* | 2042 | 1256 | 967 | 1075 | 289 | *TH:GAD* | *GAD:TH* |
|  |  |  |  |  |  |  | 1.63 | 0.62 |
|  |  |  |  |  |  |  | *A10 Total* |  |
|  |  |  |  |  |  |  | *TH:GAD* | *GAD:TH* |
|  |  |  |  |  |  |  | 1.67 | 0.60 |
| **A10 TOTAL** |  | **4,923** | **2,952** | **2,057** | **2,866** | **890** |  |  |
| Mean |  | 615 | 369 | 257 | 358 | 111 |  |  |
| SEM |  | 55 | 26 | 17 | 41 | 19 |  |  |
| REGION | CASE | Total TH | Total GAD1 | TH+GAD1 | TH alone | GAD1alone |  |  |
| SNc<br>(A9) | J38 | 2521 | 821 | 695 | 1826 | 126 |  |  |
|  | J56 | 3810 | 1615 | 1444 | 2366 | 171 |  |  |
|  | J59 | 2667 | 1225 | 1119 | 1548 | 106 |  |  |
|  | J61 | 2757 | 1106 | 1030 | 1147 | 76 |  |  |
|  | *A9 Total* |  |  |  |  |  | *TH:GAD* | *GAD:TH* |
|  |  |  |  |  |  |  | 2.47 | 0.41 |
| **A9 TOTAL** |  | **11755** | **4767** | **4288** | **6887** | **479** |  |  |
| Mean |  | 2,939 | 1,192 | 1,072 | 1,722 | 120 |  |  |
| SEM |  | 294 | 165 | 154 | 256 | 20 |  |  |
| REGION | CASE | Total TH | Total GAD1 | TH+GAD1 | TH alone | GAD1alone |  |  |
| RRF<br>(A8) | J38 | 723 | 357 | 303 | 420 | 54 |  |  |
|  | J56 | 259 | 178 | 117 | 142 | 61 |  |  |
|  | J59 | 747 | 382 | 318 | 429 | 64 |  |  |
|  | J61 | 966 | 665 | 547 | 419 | 118 |  |  |
|  | *A8Total* |  |  |  |  |  | *TH:GAD* | *GAD:TH* |
|  |  |  |  |  |  |  | 1.70 | 0.59 |
| **A8 TOTAL** |  | **2695** | **1582** | **1285** | **1410** | **297** |  |  |
| Mean |  | 674 | 396 | 321 | 353 | 74 |  |  |
| SEM |  | 149 | 101 | 88 | 70 | 15 |  |  |
|  |  |  |  |  |  |  | *Pooled Subregions Total* |  |
|  |  | **Total TH** | **Total GAD1** | **TH+GAD1** | **TH alone** | **GAD1alone** | **TH:GAD** | **GAD:TH** |
| **POOLED SUBREGION TOTALS** |  | **19,373** | **9,301** | **7,630** | **11,163** | **1,666** | 2.08 | 0.48 |
| Mean |  | 1,211 | 581 | 477 | 698 | 104 |  |  |
| SEM |  | 269 | 102 | 98 | 165 | 12 |  |  |

### Statistical analysis

Individual counts for each neuronal combination (TH-only, VGluT2-only, GAD1-only, TH+VGluT2, TH+GAD1, VGluT2+GAD1, and TH+VGluT2+GAD1) in each DA subregion were summed for each case and normalized to total counts per region to account for region size. Across animal comparisons were performed using a 2-way ANOVA (main effects only) with Tukey’s multiple comparisons test (with single pooled variance). Individual cases were analyzed separately and compared to all other cases. No significance differences were found across cases (2-way ANOVA; F(4,24)=1.9; p>0.999). Similarly, across sex comparisons (n=3 males, n=2 females) were analyzed using 2-way ANOVA (main effects only) with Tukey’s multiple comparisons test (with single pooled variance) and found to be insignificant (F(1,6)=3.02, p=0.9987). We therefore pooled the data for each individual subregion (regardless of or case) for both the RNAscope transcript and ICC protein immunoreactivity experiments. Combined region and individual DA subregion comparisons were quantified following a two-way ANOVA with Tukey multiple comparisons test (corrected) to assess total counts across neuronal combination categories. p<0.05= ^*^, p<0.01=^**^, p,0.001= ^***^, p<0.0001=^****^. Error bars are presented as SEM.

## Results

### RNAscope combinatorial analysis of TH+, VGluT2+ and GAD1+ transcripts: overview

Seven neuronal phenotypes were found in the ventral midbrain in all DA subregions: TH+ only, VGluT2+ only, GAD1+ only, TH+/VGluT2/GAD1 [triple], TH+GAD1, GAD1+VGluT2 and TH+VGluT2. CaBP-positive subregions are the mVTA (outlined in black), PBP (outlined in yellow) and RRF (outlined in blue) subregions, while CaBP is absent in the SNc (outlined in pink) **(Fig 1A-E**). After tracings were overlaid onto adjacent sections processed for RNAscope (**Fig 1F-J)**, determinations if transcript expression and co-expression were made at higher magnification. Representative images in the VTA (**Fig 1K**), PBP (**Fig 1L**), SNc (**Fig 1M**) and RRF (**Fig 1L)** showed various cell-types (transcript combinations) including TH+VGluT2+GAD1 (triple, white), TH+VGluT2 (pink), VLGUT2+GAD1 (yellow), TH alone (blue), VGluT2 alone (red) and GAD1 alone (green). TH+GAD1 not shown.

**Figure 1.**
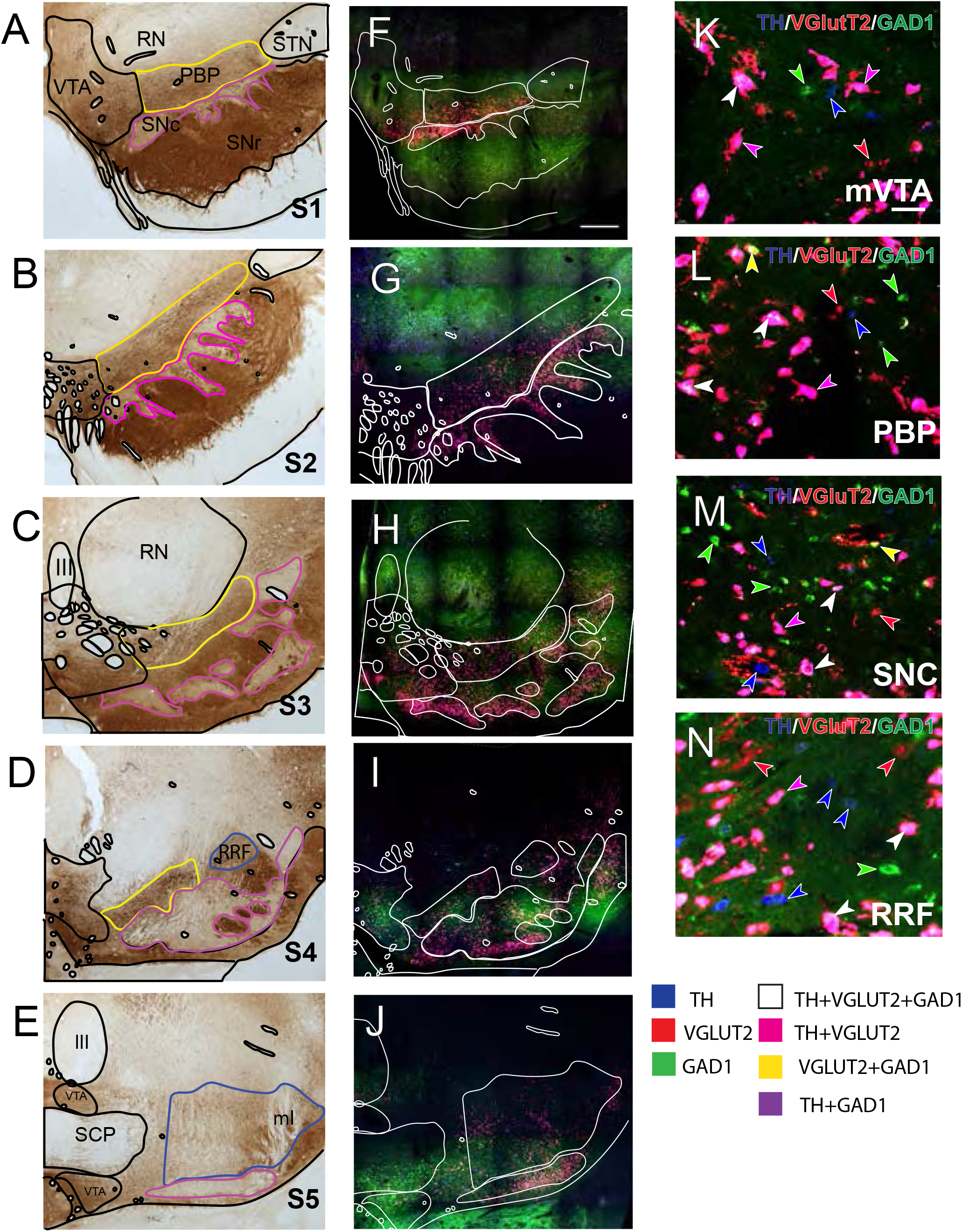
TH, VGluT2, and GAD1 transcript labeling in DA subregions. *A-E)*. Calbindin (CaBP) immunohistochemical reactivity delineates DA subregional boundaries through the rostrocaudal extent of the ventral midbrain, with DA neurons co-containing CaBP in the A10 and A8, and SNc (A9) is devoid of CaBP immunoreactivity. The midline VTA (A10, outlined in black), PBP (A10, outlined in yellow) and RRF (A8, outlined in blue) are shown. F-J). Regional boundaries are overlaid on adjacent RNAscope-processed sections with careful alignment of fiducial markers such as blood vessels and fiber tracts. K-N). Representative examples of single, double and triple transcript labeling of neurons in midline VTA (K), PBP (L), SNc (M) and RRF (N). TH-single label (blue arrow), TH-VGluT2-GAD1 labeled (white arrow), TH-VGluT2 labeled (pink arrow), and VGluT2-GAD1 labeled (yellow arrow), GAD1 single labeled (green arrow), and VGluT2-single labeled examples are shown. Scale bar= 50um. Abbreviations: GAD1, glutamate decarboxylase 1; ml, medial lemniscus; PBP, parabrachial nucleus; RRF, retrorubral field; RN, red nucleus, SCP, decussation of the superior cerebellar peduncle; SNc. substantia nigra, pars compacta; SNr, substantia nigra pars reticulata; STN, subthalamic nucleus; TH, tyrosine hydroxylase; mVTA, midline nuclei of the ventral tegmental area,; VGluT2, vesicular glutamate transporter 2; III, fascicles of the third nerve.

### Distibution of neuronal phenotypes in the ventral midbrain: labeled neurons versus single-labeled phenotypes

Prior studies, including ours, have analyzed neuron counts using single-label approaches. To better interpret these findings, we analyzed the relative distribution of cells containing TH, VGlut2, and GAD mRNA versus single labeled cells (**Fig 2A, B**, rostro-central and **2E, F** caudal). All subregions had many neurons containing TH, VGluT2 and GAD1 (**Fig 2A**, rostral; **2E**, caudal), with fewer single labeled neurons across all DA subregions. The four types of multiplex neuronal subpopulations (TH+VGluT2+GAD1 [triple], TH+GAD1, GAD1+VGluT2, TH+VGluT2) were represented in all DA subregions, higher concentrations of TH+VGluT2+GAD1 [triple] and TH+VGluT2 phenotypes predominating.

**Figure 2.**
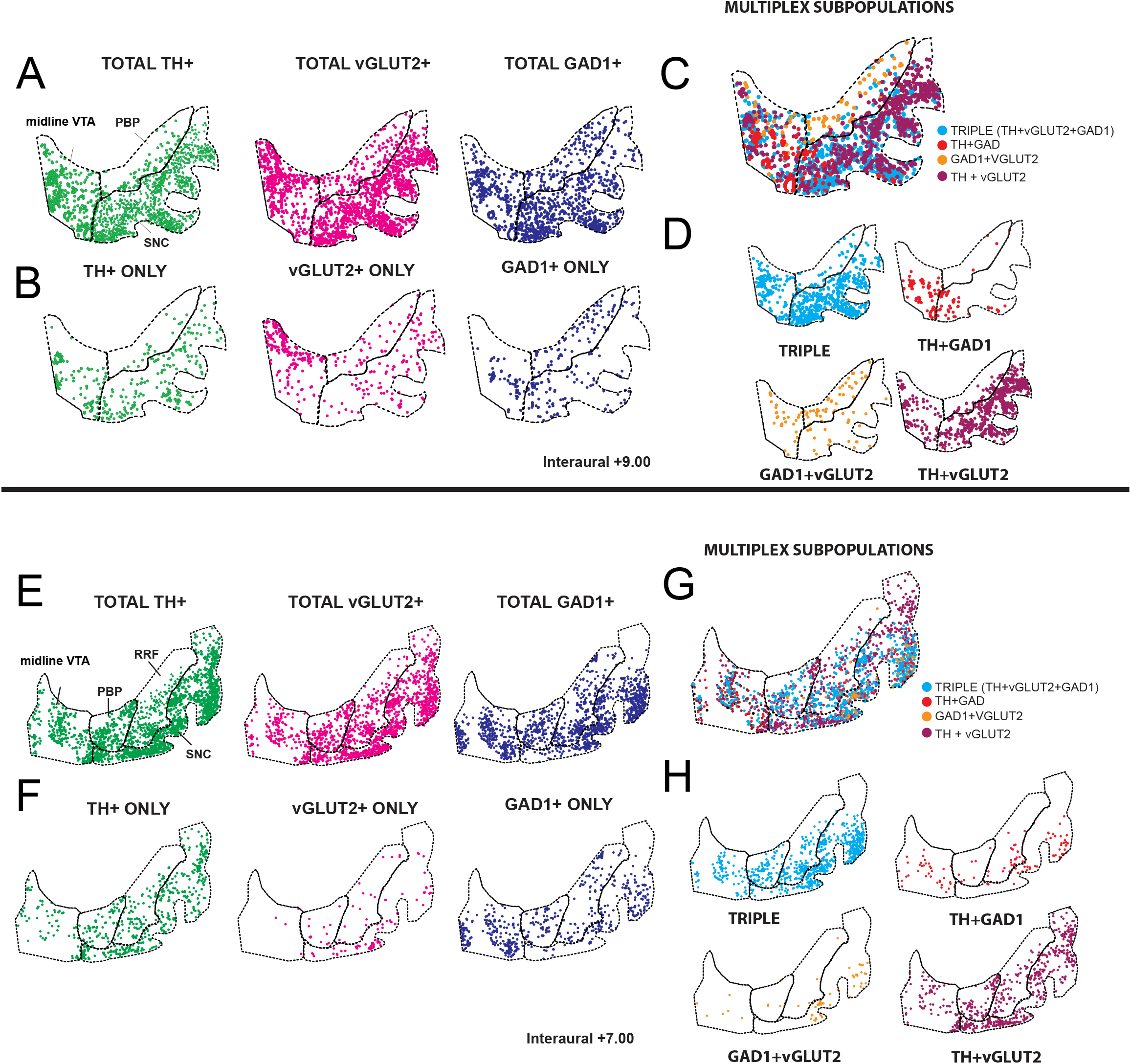
Mapping of single and multiplexed neurons in the ventral midbrain. A-B and E-F). Comparison of the distribution of all neurons containing one of the three transcripts (TH, green; VGluT2, red; GAD1, blue) versus the population of single-labeled neurons in the rostral (A-B) and caudal (E-F) midbrain. C-D and G-H). Multiplex neuron neurons only at rostral (C-D) and caudal (G-H) levels of the ventral midbrain. All neuronal phenotypes are found across all DA subregions (dashed line).

#### Quantification of neuronal phenotypes with mRNA

Consistent with the impression from qualitative maps in Fig. 2, normalized counts (# individual phenotype/total # all phenotypes) revealed that TH-VGlut2 (22%) and TH-VGlut2-GAD1 (23%) labeled neurons made up the majority of all labeled neurons (45%)(**Fig. 3A, Table 2**). TH-only and GAD1-only neurons had lower proportions(19%, 20% respectively), and VGluT2 - alone neurons had the lowest proportion (8**%**).

**Figure 3.**
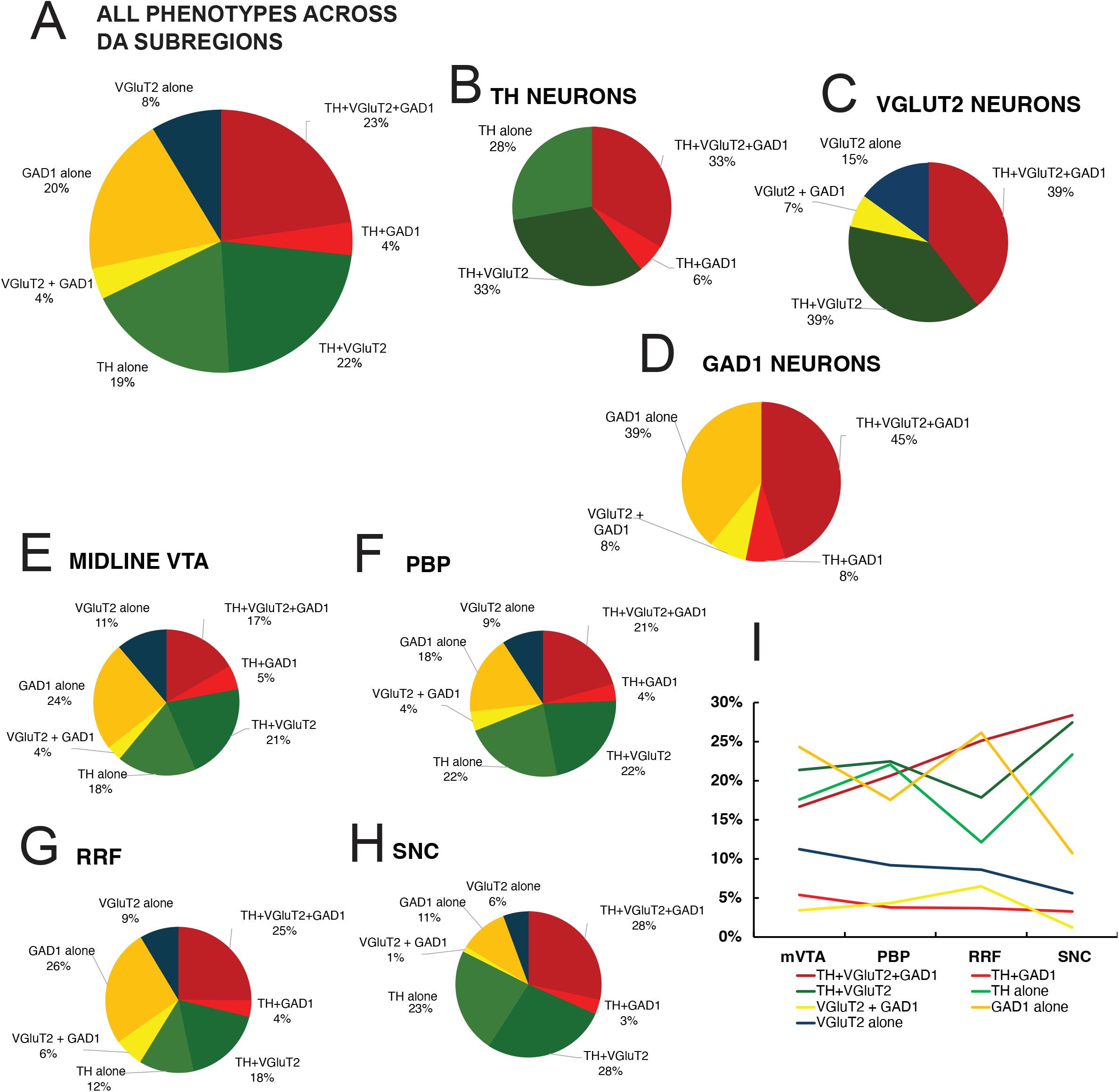
Quantification of RNAscope data. A) Proportions of all 7 neuronal phenotypes across all subregions for all animals. B) Proportion of TH mRNA + neurons that are single-labeled, dual- and triple-labeled. C) Proportion of VGluT2-containing neurons that are single, dual, and triple-labeled. D) Proportion of GAD1+ mRNA neurons that are single, dual, and triple-labeled. E-H). Percentage of single/multiplex neuron types by individual DA subregion. I.) Relative frequency histogram of the 7 phenotypes in each DA subregion. TH-only, TH+VGLUT2+, and TH+VGLUT2+GAD1+ consistently displayed the highest incidence of expression. TH+GAD1+ and VGLUT2+GAD1+ cell types remained the lowest combinations expressed in all regions.

Considering all TH+ cells as a group, it was notable that TH mRNA was expressed predominantly with VGluT2 (66%) (**Fig 3B**). Overall, TH mRNA was expressed with another transcript 72% of the time, indicating that co-expression is the rule rather than the exception among DA neurons as a group. Interestingly, while TH-VGlut2-GAD1+ neurons were abundant (33%), TH mRNA was infrequently found with GAD1 mRNA alone (6%). This finding appears to link GAD1 expression with VGlut2 expression in TH-expressing neurons.

Among all neurons expressing VGluT2 mRNA, VGluT2 mRNA was expressed with one or more transcripts 85% of the time, with TH as the main co-transcript the majority of the time (**Fig 3C**, 78%). VGlut2-single labeled cells formed only 15% of all VGlut2-containing neurons. In GAD1 mRNA labeled neurons, GAD1 was mostly co-expressed with both TH and VGluT2 (53%, **Fig 3D**,) with GAD1+-only expressing cells (presumptive interneurons) making up the next largest population (39%). VGluT2-GAD1(8%) and TH-GAD1 phenotypes (8%) were least represented.

We next sought to determine the neuronal phenotypes across DA subregions (**Fig 3E-H**). Surprisingly, the proportions of each phenotype within each DA subregion mirrored those found in grouped data (**Fig 3A**). The largest contribution was by multiplexed TH+ neurons (TH+VGluT2; 18-28%, and TH+ VGluT2+GAD1+ (17-28)). To determine whether different DA subregions displayed preferential neuronal combinations, we did a multi-comparisons analysis comparing the neuronal combination type across different DA subregion. Across DA subregions, the mean proportion of each neuronal phenotypes was similar (**Fig 3E-H**), with one exception. TH+VGluT2+GAD1+ neurons were higher in SNc compared to other subregions, reaching significance in the mVTA comparison where only 17% TH+VGluT2+GAD1 neurons were found (one-way ANOVA, F[1.44, 5.8]=9.87, p=0.017; Tukey’s multiple comparisons test, p=0.0041). Summing TH+ neurons of all phenotypes revealed that they were 1.5-1.9x higher than GAD1 + single-labeled neurons in mVTA, PBP and SNc-, compared to the RRF where the ratio was 1.2x, consistent with previous stereologic results ^37^.

### TH and GAD1 protein expression and co-expression correlates with mRNA transcript distribution

A primary function of mRNA is to act as a template for protein biosynthesis. We therefore next sought to investigate the expression patterns of TH and GAD1 to determine if the pattern of neuronal phenotypes for TH and GAD1 was retained at the protein level. We then extrapolated results to account for VGlut2 expression which could not be directly measured. Immunohistochemical processing revealed many brightly single-labeled TH+ and GAD1+ cells that were easily discernable based on morphology (**Fig 4A-E**). TH+ neurons (green) were heterogeneous, and often notably large with elongated cell bodies with thick proximal branches extending out to thinner processes, as previously described^7, 37^(**Fig. 4C-D**). Many of these had GAD1+ colocalization. Single-labeled GAD1+ neurons were often smaller and round. Qualitatively, we noted these three neuronal phenotypes in all DA subregions throughout the rostral to caudal extent of the ventral midbrain (**Fig 4F-H**). TH+ neurons and TH-GAD1+ neurons were densest in the SNc as expected. GAD1+ single labeled neurons were a smaller population overall, that was more densely distributed in the A10 (midline VTA and PBP) and A8 subregions.

**Figure 4.**
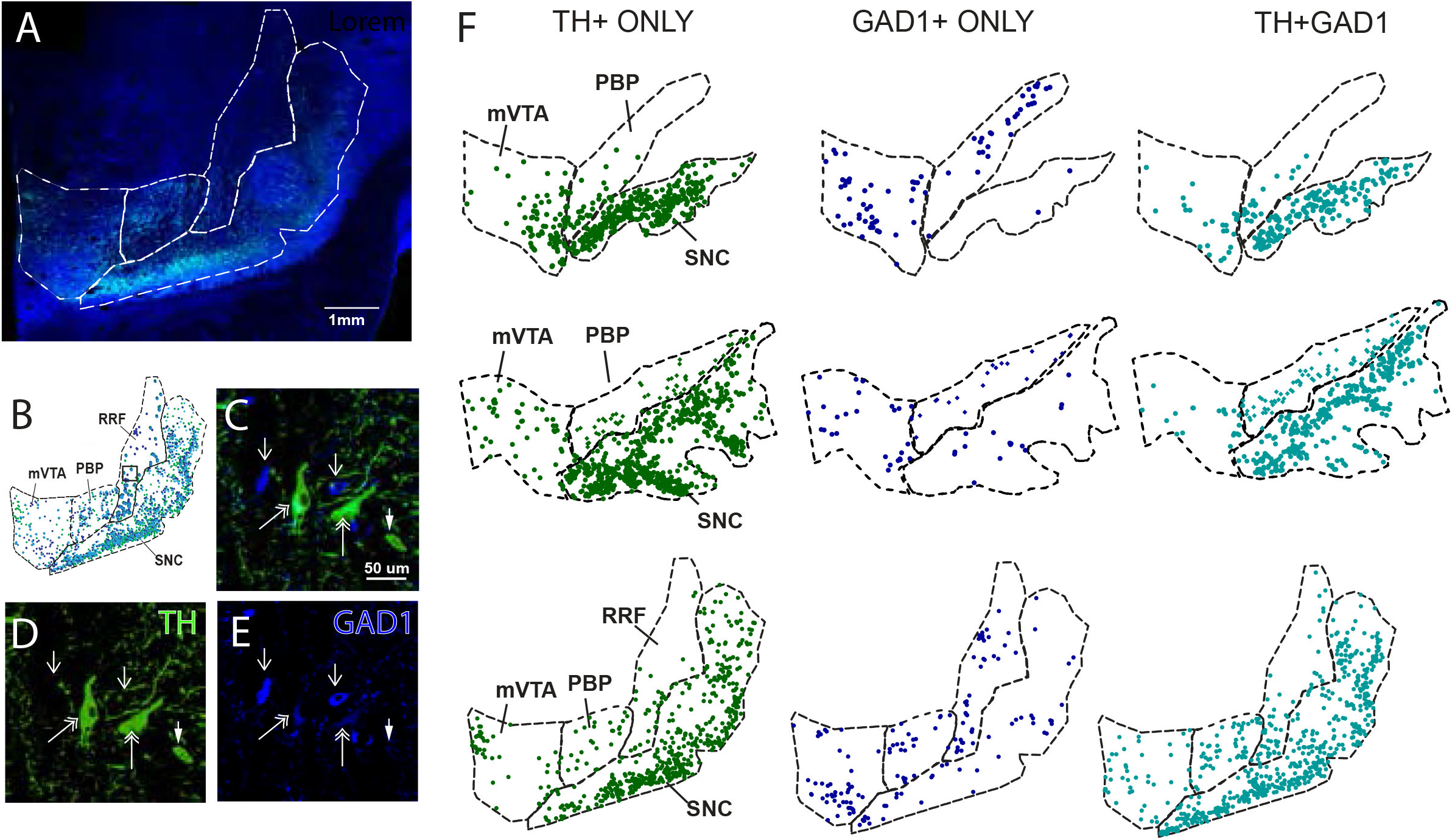
Mapping TH+, GAD1+, and TH+GAD1+ immunoreactive neurons across the ventral midbrain. A). Macroscopic view of a mid-caudal section immunoreacted for TH (dopaminergic neurons, green) and GAD1 (GABAergic neurons, blue). DA subregions are depicted with dashed white lines. B.) Representative plot (image in A) after colocalization, with TH+-only (green dots), GAD1+-only (blue dots) neuron, and TH+GAD1+ dual labeled neurons (cyan dots). C-E.) A magnified photomicrograph, with TH in the green channel, GAD1 in the blue channel, and double labeled neurons in cyan. Single-labeled TH+ neurons (green, thin, single arrowhead), single-labeled GAD1+ neurons (blue, solid arrowhead; AlexaFluor 647), and TH+GAD1+ overlap (cyan, thin, double arrowhead) are shown. Scale bar-50 um. F.) Representative maps of TH+ only (green), GAD1+ only (blue), and TH+GAD+ (cyan) neurons in rostral-caudal sections (top to bottom).

We quantified the total neuron counts for TH+, GAD1+, and TH+GAD1+ in all DA subdivisions (midline VTA, PBP, RRF and SNc) including all sections through the rostral to caudal axis (**Fig 5, Table 3**). Overall, dual labeled TH+GAD1 (37%) formed a large subset of TH-containing neurons, although TH+ only neurons were the largest population. GAD1-only neurons comprised a relatively small percentage of all labeled neurons (8%). There was a significant effect between neuronal phenotype and cell counts (one-way ANOVA, F[2,29]=89.23, p<0.0001) as well as significant between group comparisons (**Fig 5B**, Tukey’s Multiple Comparisons Test; TH+GAD1 vs TH-only, p=0.002; TH+GAD1 vs GAD1-only, p<0.0001; TH-only vs GAD1-only, p<0.0001). Similar differences continued among the three neuronal phenotype counts for each DA subregion (**Fig 5C-F**). Within the midline VTA (**Fig 5C**), GAD1-only labeled cells had the highest percentage of all subregions (F[1,3.2]=38.78, p=0.0066), and was not significantly from TH+GAD (p=0.1106). Significant differences were found between TH+GAD and TH-only (p=0.0008), and TH-only and GAD1-only, phenotypes (p=0.0121). In PBP (**Fig 5D**), TH+GAD1 and TH-only groups were each significantly higher than GAD1-only labeled cells (p=0.0087 and 0.0142, respectively), but not significantly different from each other (p=0.5367).

**Figure 5.**
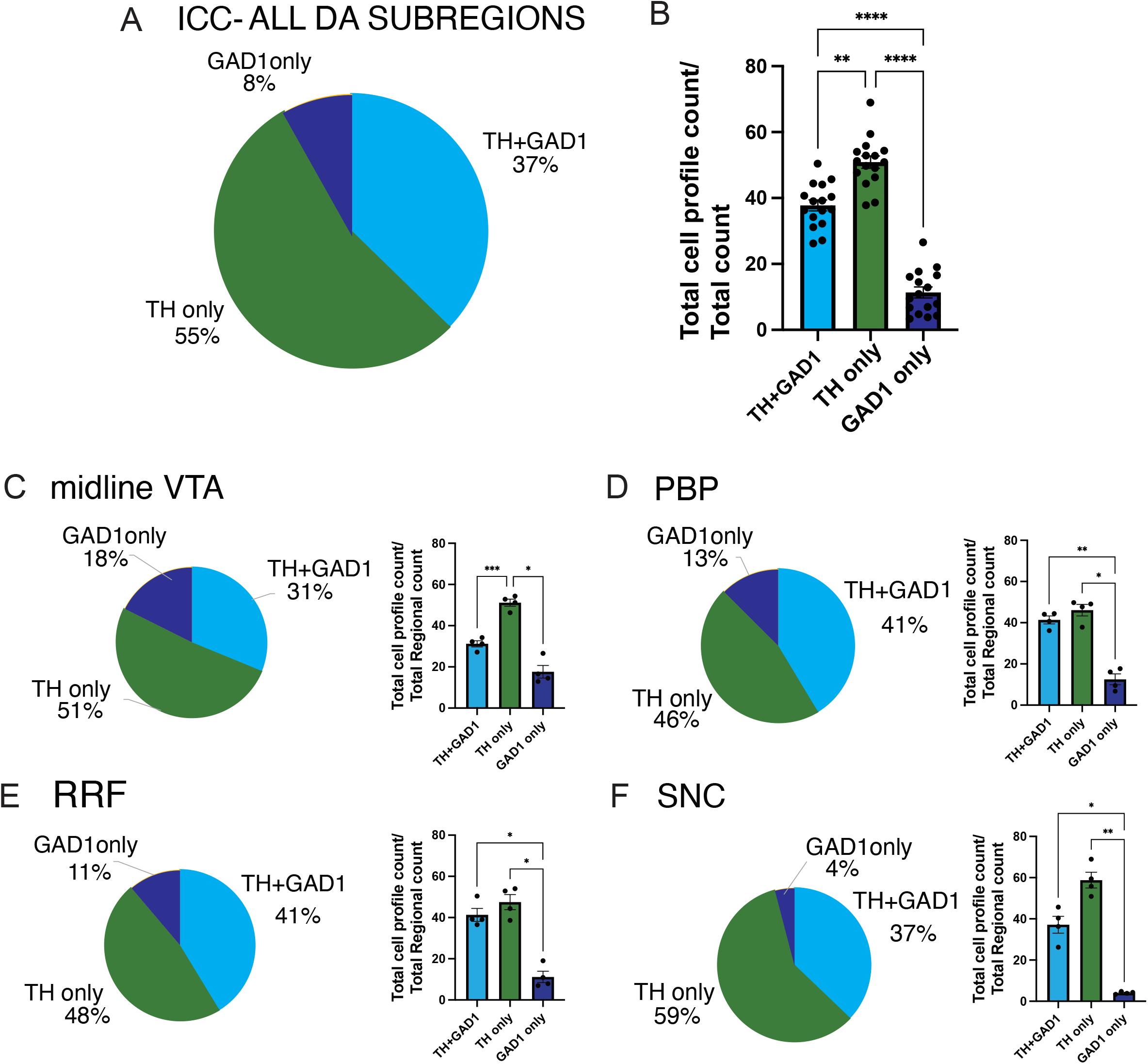
Quantification of ICC data. A) Proportions of the three phenotypes (TH-only, GAD1-only, and TH-GAD1) in all DA subregions. (B) Histogram of normalized counts. C-F) Percentage ICC cell profile combinations in individual in midline VTA (C), PBP (D), RRF (E), and SNc (F) with accompanying histograms. p<0.05= ^*^, p<0.01=^**^, p,0.001= ^***^, p<0.0001=^****^. Error bars are presented as SEM.

There was a significant overall difference across cell types (F[1.7,5.1]=35.97, p=0.001). The pattern in the RRF (**Fig 5E**) was very similar to that in the PBP, with TH+GAD1 and TH-only groups were each significantly higher than GAD1-only cell types (p=0.0148 and 0.0157, respectively), and no significant effect between TH-only and TH-GAD+ populations (p=0.6270). There was also an overall significant effect for all comparisons (F[1.8,5.4]=24.86, p=0.0021). The SNc also followed the pattern for the PBP, and RRF (**Fig 5F**). TH-only and TH+GAD1 groups were each significantly greater than GAD1-alone (p=0.01 and p=0.0012, respectively), with no significant effects between the TH-alone and TH-GAD+ cell groups (p=0.1397), and an overall significant effect for all comparisons (F[1,3]=48.06, p=0.0061). Analyzing the data by rostro-caudal trajectory (approximately 6 mm), TH+ cell groups (TH+ green; TH-GAD1+ cyan) vastly outnumbered GAD1+ cells (yellow) throughout this entire trajectory, gradually increasing until level S4 (**Suppl Fig 1A-E**). GAD1+-only cell counts relatively low throughout all levels. Subregional patterns were largely similar to this overall trajectory.

To compare the mRNA and protein expression, we normalized the proportion of mRNA phenotypes to the total population, subtracting out VGluT2 phenotypes. The proportion of TH+-only, TH+/GAD1+, and GAD1+ neurons in the protein expressing study (absent glutamate markers) was then compared to the mRNA phenotypes (**Fig. 6**). The proportion of TH+ only neurons in the immunohistochemistry study was 55% compared to 51% TH+ neurons in the mRNA study (including TH+ only neurons and TH+-VGluT2 phenotypes). TH-GAD1 neurons in the immunocytochemistry study were 37% versus a total of 30% TH-GAD1 phenotypes (TH-VGluT2-GAD1 plus TH-GAD1 subpopulations) in the mRNA study, suggesting relatively similar correspondence between TH transcript and protein levels. The greatest mismatch was for GAD1+ neurons lacking TH, which were 8% in the protein expression study, versus 19% in the mRNA study, accounting for GAD1-alone and VGluT2-GAD1 phenotypes. GAD1 mRNA expression and translation dynamics may explain this mismatch, perhaps reflecting delayed translation of transcripts and/or slowed degradation of transcripts after translation in one or both subpopulations. There is frequently not a linear relationship between transcript and protein expression, depending on cell type ^45^, which cannot be assessed in our data.

**Figure 6.**
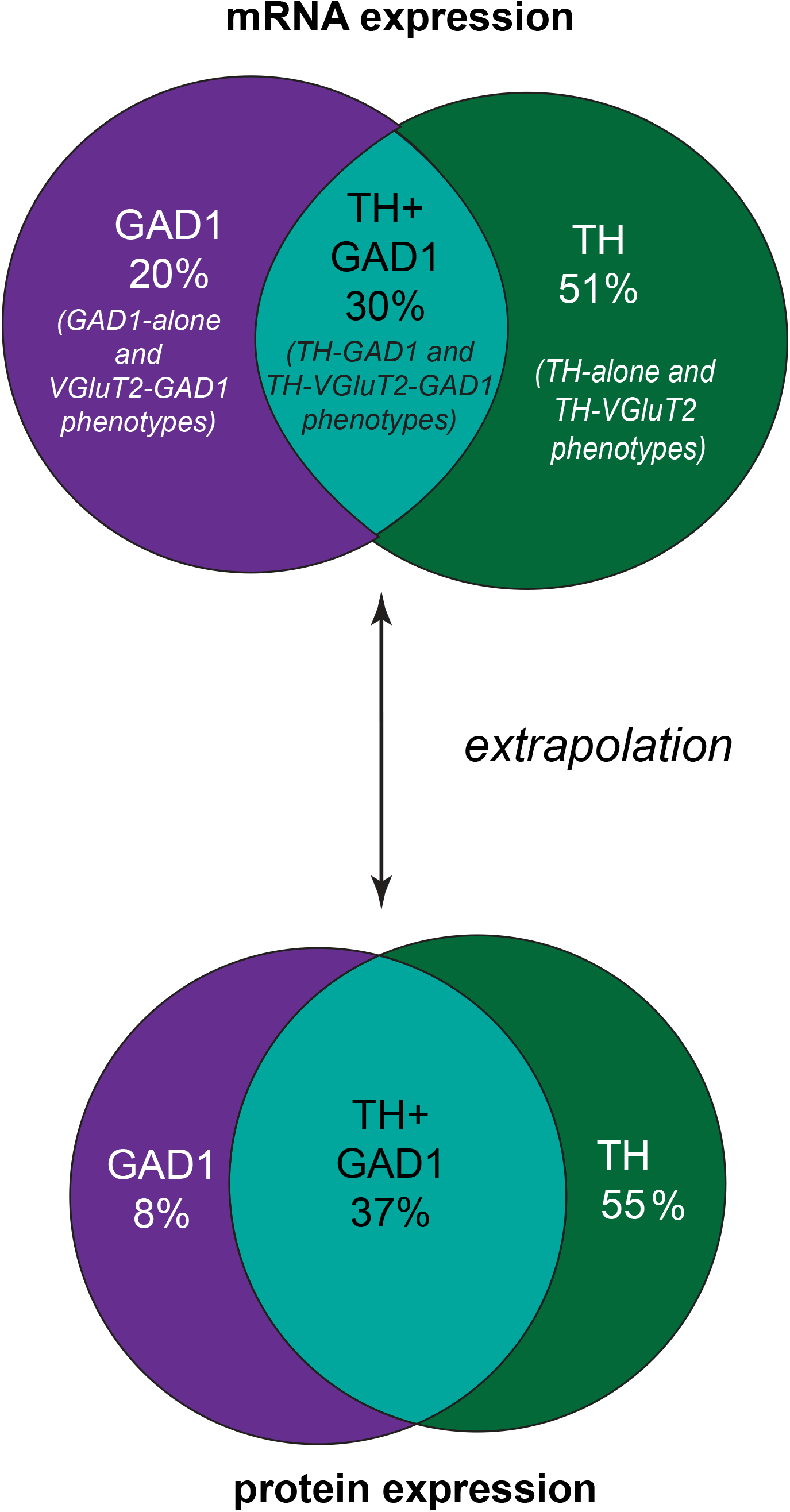
Comparison of RNAscope and ICC phenotypes. Extrapolation of neuronal expression patterns (see Methods) between mRNA and protein expression data. *(VGluT2 single-labeled neurons are subtracted from the denominator for mRNA percentages)*.

## Discussion

In young macaques, the RNAscope technique yielded robust TH, VGluT2 and GAD1 transcript expression throughout the ventral midbrain, yielding 7 different phenotypes. The most prevalent cell types were multiplexed DA neurons, followed next by TH-only DA neurons. GAD1 and VGluT2 frequently were mostly found as likely co-transmitters. Surprisingly there was little difference in the relative expression of cell types across DA subregions. These finding were reinforced following protein determinations where we found that mRNA and protein expression followed similar--although not exact--patterns suggesting that mRNA is largely manufacturing functionally relevant protein.

### Species, age, and methodologic considerations: the midbrain in adolescent macaques

In this cohort, there is a larger proportion and broader distribution of multiplexed DA neurons in the ventral midbrain, compared to many prior reports in rodents, marmosets and human. Most notably, 68% of all TH+ neurons contained another neurotransmitter marker, with 45% of these neurons containing VGlut2 mRNA. While we anticipated robust expression of TH, the finding that a large percentage were ‘multiplexed’ was surprising, based on work in other species and in older primates ^28, 29, 46-49^. Although VGluT2-dependent glutamate release by dopamine neurons has been described for 20 years, the anatomical expression patterns of VGluT2 mRNA in TH-containing cells in rodent models is consistently relatively low, and largely limited to the VTA ^24^(Review ^50^).

Discrepancies across studies may lie in many factors, including methodological approaches. In mice, in situ hybridization studies indicates variable but generally low levels of VGluT2 expression by dopamine neurons ^47, 51, 52^. However, in other reports using single cell RT-PCR, the proportion of DA neurons containing VGluT2 mRNA appear significantly higher ^53^, such that it has been concluded that standard *in-situ* hybridization techniques may not be sufficient to resolve VGluT2 mRNA transcript reliably ^50, 54^. Further, the application of immunohistochemistry to detect TH signal following ISH for VGluT2 ^29, 47, 49^ introduces the potential of masking or diminishing transcript signal. Indeed, the advent of RNAscope, a more sensitive branched in-situ hybridization technique, provides the ability to detect mRNA transcript visualization in thicker sections due to an amplified fluorescent signal ^55^, thus resolving potential populations of neurons that were previously overlooked. Some later reports using genetic approaches ^56-58^ and/or RNAscope approaches have found slightly increased expression of multiplexed neurons in the VTA in mice ^26, 46, 51^.

Another important consideration for VGluT2 expression is the age of the animal, which has not been consistently considered ^59-61^. Developmental comparisons of mouse versus human brain at different ages found age-related decreases in TH and VGluT2 mRNA expression from adolescent to middle age to senescence in both species ^61^, supporting earlier work showing that young animals have higher levels of both TH- and VGluT2-containing neurons ^59, 62, 63^. TH+ neurons (detected by ICC) in young (3-5 year old, adolescent) macaques, are twice as numerous as in old animals (22-25 years) ^64^. The animals in our study fall within a juvenile-adolescent age group (3-4 years) and ‘young adult’ time-points (6.5 years)^38^, and exhibited robust TH-labeling, with the majority of TH+ neurons co-containing VGluT2 mRNA (66%). Consistent with the robust signal for TH+/VGluT2+ neurons across the midbrain in our young animals, in human, sampling TH+/VGluT2+ mRNA neurons in both the VTA and SNc revealed highest counts in the 16-21 year old group compared to middle-age and elderly group ^61^.

### GAD1 and VGluT2 co-expression in DA neurons

GABA (detected by GAD1) and glutamate (detected by VGluT2) appear to function primarily as co-transmitters with TH--lending flexibility to transmitter release properties at the terminal ^27^. These fast-transmitter associated transcripts were co-expressed predominantly with TH mRNA (78% of all VGluT2+ neurons; 53% of all GAD1+ neurons), suggesting complexity in dopaminergic signaling in young animals that is only beginning to be explored in the primate brain.

Over a decade of electrophysiologic evidence in rodents shows that DA neurons mediate a fast excitatory signal ^24, 65-68^. Yet anatomic identification of neurotransmitter co-localization in midbrain neurons is challenging, not only for the technical reasons noted above, but also because the typical markers that stand in for transmitters are enzymes or vesicular markers, with shifting production or function over development or stress. Although TH is a widely used marker for DA in the midbrain ^2^, its mRNA and protein can be up- and down-regulated with normal aging ^61, 64^ or stress^69, 70^. Controlling for age and stress environment (see ^39^), we found TH mRNA and protein markers to be highly stable across animals. The dopamine transporter (DAT), which governs DA reuptake, and the vesicular monoamine transporter 2 (VMAT2), are other common indicators of DA neurons ^71^. However, in monkey, DAT has different expression levels in specific DA subregions ^35^, and VMAT mRNA has broader packaging properties beyond DA, e.g. of GABA ^25, 72^.

Detection of GAD1 mRNA in a TH-VGluT2 subpopulation was surprising, as it has been notably absent from DA neurons in several assays in mice, prompting alternate theories for GABA production ^73^, and/or re-utilization at the synapse ^74^. On the other hand, GAD2 mRNA is co-localized with dopamine markers in restricted cell populations ^75, 76^. A key route for GABA synthesis, GAD1 is an enzyme that makes use of recycled glutamate, by converting glutamine to GABA. Interestingly, we found that GAD1mRNA co-expression in TH+ neurons was almost always in TH+VGluT2+mRNA phenotypes. In contrast, there were relatively few TH neurons co-expressing GAD1 mRNA without VGluT2, suggesting the possibility that GABA may be synthesized from glutamine in the triple-labeled DA phenotype.

### Functional-Development Relevance of Multiplex DA Neurons in Young Macaques

Given the wide role DA plays in a variety of fundamental behaviors, it is unsurprising that transmitter signaling mechanisms of DA neurons would also be complex ^77^. The multiplex phenotype may have different release mechanisms, depending on whether neurotransmitters are co-packaged in the same synaptic vesicle (co-release) or in separate synaptic vesicle populations (co-release or co-transmission)(see review by Eskenazi ^77^). In addition to having different pre- and post-synaptic receptor effects, co-leased transmitters can modulate the rate of packaging of transmitters into synaptic vesicles^78^.

At the circuit level, studies in adult mice link specific transmitter phenotypes to specific DA regions and circuits (Review, Poulin ^79^). For example, DA-VGluT2 containing neurons in the midline VTA nuclei, project to the shell of the nucleus accumbens ^20, 52, 80, 81^. Fewer DA-VGluT2 neurons in the lateral VTA or SNc project to the nucleus accumbens core, dorsal striatum, and tail of the striatum^22, 82^. A small number of DA-VGluT2 neurons project to the rodent prefrontal cortex (PFC), usually limited to the infralimbic and prelimbic cortex^83-85^ (however, see ^86^). This differential distribution of DA neuron circuits appears to have differential physiologic effects on striatal subregions—and potentially the PFC. It is hypothesized that glutamatergic physiologic responses at DA terminals have important implications for synaptic plasticity ^22, 67, 68^.

In the nonhuman primate, DA projections to the striatum form a gradual step-wise pattern, and not strictly divided by the VTA and SNc as in rodents ^87, 88^. Moreover, DA inputs to the cortex are widespread covering much of the cortical mantel ^89, 90^, and emanating from the entire mediolateral sweep from the A10 (including the PBP) through the A8 ^90-92^. In monkey, DA neuron responses to appetitive and aversive stimuli vary along this trajectory ^93^. Here, the widespread distribution of all DA phenotypes across all DA subregions, suggests that mixtures of DA phenotypes target the entire forebrain, at least in young animals.

Development plays a critical role in DA neuron phenotypes, which begins in the embryo but continues postnatally. Although there are basic similarities in DA neuron gene expression shared in mouse and human, there are also many dissimilarities, with final phenotypes developing postnatally ^76, 94^. Perhaps the largest variable in DA phenotype is developmental timing, which in long-lived species may occur over many years. In rodent species, all DA-destined neurons are initially glutamatergic, with many fewer DA-glutamatergic neurons by adolescence and into adulthood ^59, 60, 63^. Glutamate action on DA neurons is related to axonal lengthening, arborization, and survival ^95, 96^, and is implicated in synapse formation in young animals^60, 97, 98^. Interestingly, the meso-cortical projection is late-developing, and extends through the adolescent period across species (see Reviews, Reynolds^99^ and Islam^100^). While speculative, the abundant DA-VGluT2 population in our young animals, may represent a mechanism for the normative axon extension governing this pathway in adolescence.

## Conclusion

In summary, there has been considerable progress made in defining the anatomical distribution of dopaminergic cell types in the ventral midbrain. Here we report the distribution of 7 distinct NT profiles in the macaque midbrain utilizing high-resolution RNAscope and ICC confirmation. Our findings of robust levels of multiplex neurons throughout the 3 primary subregions of the ventral midbrain (A10, A9, and A8) may be the result of advanced methodologies, primate model, and/or age of subjects. Our data support and expand on earlier work demonstrating heterogeneity within the dopaminergic cell regions in rodents, contributing to complex downstream dopaminergic circuitry, diverse signaling and function. Our work also aligns with emerging data on age-dependent changes in DA phenotypes in higher species.

## Acknowledgements

We thank the Center for Advanced Light Microscopy and Nanoscopy at the University of Rochester Medical Center for guidance and advice. Dr. Mahoui is now in the Department of Family Medicine, University of California, Davis.

## FIGURE LEGENDS

Supplemental Figure 1. Quantification of TH+GAD1, TH-only, and GAD1 only expressing neurons across the rostral to caudal axis. A.) TH-only (green line), TH+GAD1 (red line), and GAD1 (yellow line) were consistent in their relationship to one another across the rostrocaudal axis. Average cell counts for each cell-type at levels S1-6 is shown. B-D.) Rostrocaudal patterns, compared across DA subregions, were similar. The majority of TH-containing neurons is in the SNc, as expected.

## References

1. Pearson J, Goldstein M, Brandeis L. Tyrosine hydroxylase immunohistochemistry in human brain. Brain Res 1979; 165: 333–337.

2. Kitahama K, Sakamoto N, Jouvet A, Nagatsu I, Pearson J. Dopamine-beta-hydroxylase and tyrosine hydroxylase immunoreactive neurons in the human brainstem. Journal of chemical neuroanatomy 1996; 10(2): 137–146.

3. Arsenault MY, Parent A, Seguela P, Descarries L. Distribution and morphological characteristics of dopamine-immunoreactive neurons in the midbrain of the squirrel monkey (Saimiri sciureus). J Comp Neurol 1988; 267: 489–506.

4. Beier KT, Steinberg EE, DeLoach KE, Xie S, Miyamichi K, Schwarz L et al. Circuit Architecture of VTA Dopamine Neurons Revealed by Systematic Input-Output Mapping. Cell 2015; 162(3): 622–634.

5. Bjorklund A, Dunnett SB. Dopamine neuron systems in the brain: an update. Trends in neurosciences 2007; 30(5): 194–202.

6. Morales M, Margolis EB. Ventral tegmental area: cellular heterogeneity, connectivity and behaviour. Nature reviews Neuroscience 2017; 18(2): 73–85.

7. Pearson J, Goldstein M, Markey K, Brandeis L. Human brainstem catecholamine neuronal anatomy as indicated by immunocytochemistry with antibodies to tyrosine hydroxylase. Neuroscience 1983; 8, No. 1: 3–32.

8. Watabe-Uchida M, Zhu L, Ogawa SK, Vamanrao A, Uchida N. Whole-brain mapping of direct inputs to midbrain dopamine neurons. Neuron 2012; 74(5): 858–873.

9. Zell V, Steinkellner T, Hollon NG, Warlow SM, Souter E, Faget L et al. VTA Glutamate Neuron Activity Drives Positive Reinforcement Absent Dopamine Co-release. Neuron 2020; 107(5): 864–873 e864.

10. Yoo JH, Zell V, Gutierrez-Reed N, Wu J, Ressler R, Shenasa MA et al. Ventral tegmental area glutamate neurons co-release GABA and promote positive reinforcement. Nature communications 2016; 7: 13697.

11. Pignatelli M, Bonci A. Role of Dopamine Neurons in Reward and Aversion: A Synaptic Plasticity Perspective. Neuron 2015; 86(5): 1145–1157.

12. Morris G, Nevet A, Arkadir D, Vaadia E, Bergman H. Midbrain dopamine neurons encode decisions for future action. Nature neuroscience 2006; 9(8): 1057–1063.

13. Murphy BL, Arnsten AFT, Jentsch JD, Roth RH. Dopamine and spatial working memory in rats and monkeys: Pharmacological reversal of stress-induced impairment. J Neurosci 1996; 16: 7768–7775.

14. Sesack SR, Grace AA. Cortico-Basal Ganglia reward network: microcircuitry. Neuropsychopharmacology : official publication of the American College of Neuropsychopharmacology 2010; 35(1): 27–47.

15. Hegarty SV, Sullivan AM, O’Keeffe GW. Midbrain dopaminergic neurons: a review of the molecular circuitry that regulates their development. Dev Biol 2013; 379(2): 123–138.

16. Berridge KC, Robinson TE. What is the role of dopamine in reward: hedonic impact, reward learning, or incentive salience? Brain Research - Brain Research Reviews 1998; 28(3): 309–369.

17. Romo R, Shultz W. Dopamine neurons of the monkey midbrain: Contingencies of responses to active touch during self-initiated arm movements. J Neurophysiol 1990; 63: 592–590.

18. Schultz W, Romo R. Dopamine neurons of the monkey midbrain: contingencies of responses to stimuli eliciting immediate behavioral reactions. Journal of neurophysiology 1990; 63(3): 607–624.

19. Bromberg-Martin ES, Matsumoto M, Hikosaka O. Dopamine in motivational control: rewarding, aversive, and alerting. Neuron 2010; 68(5): 815–834.

20. Azcorra M, Gaertner Z, Davidson C, He Q, Kim H, Nagappan S et al. Unique functional responses differentially map onto genetic subtypes of dopamine neurons. Nature neuroscience 2023; 26(10): 1762–1774.

21. Skupio U, Harris AZ, Polter AM. Untangling the multifaceted VTA responses to stress. Trends in neurosciences 2025; 48(8): 582–593.

22. Chuhma N, Mingote S, Moore H, Rayport S. Dopamine neurons control striatal cholinergic neurons via regionally heterogeneous dopamine and glutamate signaling. Neuron 2014; 81(4): 901–912.

23. Zhang S, Qi J, Li X, Wang HL, Britt JP, Hoffman AF et al. Dopaminergic and glutamatergic microdomains in a subset of rodent mesoaccumbens axons. Nature neuroscience 2015; 18(3): 386–392.

24. Hnasko TS, Chuhma N, Zhang H, Goh GY, Sulzer D, Palmiter RD et al. Vesicular glutamate transport promotes dopamine storage and glutamate corelease in vivo. Neuron 2010; 65(5): 643–656.

25. Tritsch NX, Ding JB, Sabatini BL. Dopaminergic neurons inhibit striatal output through non-canonical release of GABA. Nature 2012; 490(7419): 262–266.

26. Root DH, Zhang S, Barker DJ, Miranda-Barrientos J, Liu B, Wang HL et al. Selective Brain Distribution and Distinctive Synaptic Architecture of Dual Glutamatergic-GABAergic Neurons. Cell reports 2018; 23(12): 3465–3479.

27. Vaaga CE, Borisovska M, Westbrook GL. Dual-transmitter neurons: functional implications of co-release and cotransmission. Curr Opin Neurobiol 2014; 29: 25–32.

28. Steinkellner T, Conrad WS, Kovacs I, Rissman RA, Lee EB, Trojanowski JQ et al. Dopamine neurons exhibit emergent glutamatergic identity in Parkinson’s disease. Brain : a journal of neurology 2022; 145(3): 879–886.

29. Root DH, Wang HL, Liu B, Barker DJ, Mod L, Szocsics P et al. Glutamate neurons are intermixed with midbrain dopamine neurons in nonhuman primates and humans. Scientific reports 2016; 6.

30. Fu Y, Paxinos G, Watson C, Halliday GM. The substantia nigra and ventral tegmental dopaminergic neurons from development to degeneration. Journal of chemical neuroanatomy 2016; 76(Pt B): 98–107.

31. Haber SN, Fudge JL. The primate substantia nigra and VTA: Integrative circuitry and function. Crit Rev Neurobiol 1997; 11(4): 323–342.

32. Halliday GM, Tork I. Comparative anatomy of the ventromedial mesencephalic tegmentum in the rat, cat, monkey and human. J Comp Neurol 1986; 252: 423–445.

33. Yamada T, McGeer PL, Baimbridge KG, McGeer EG. Relative sparing in parkinson’s disease of substantia nigra dopamine neurons containing calbindin-D28K. Brain Res 1990; 526: 303–307.

34. Lavoie B, Parent A. Dopaminergic neurons expressing calbindin in normal and parkinsonian monkeys. Neuroreport 1991; 2, No. 10: 601–604.

35. Haber SN, Ryoo H, Cox C, Lu W. Subsets of midbrain dopaminergic neurons in monkeys are distinguished by different levels of mRNA for the dopamine transporter: Comparison with the mRNA for the D2 receptor, tyrosine hydroxylase and calbindin immunoreactivity. J Comp Neurol 1995; 362: 400–410.

36. McRitchie DA, Hardman CD, Halliday GM. Cytoarchitectural distribution of calcium binding proteins in midbrain dopaminergic regions of rats and humans. Journal of Comparative Neurology 1996; 364(1): 121–150.

37. Kelly EA, Contraras JM, Duan A, Vassell R, Fudge JL. Unbiased stereological estimates of dopaminergic and GABAergic neurons in the A10, A9 and A8 subpopulations in the young male Macaque. Neuroscience 2022; 496(496): 152–164.

38. Plant TM, Terasawa, Ei, Witchel, Selma Feldman. Puberty in Non-human Primates and Man. 5 Physiological Control Systems and Governing Gonadal Function 2015; Fourth Edition: 1487–1536.

39. Kelly EA, Love TM, Fudge JL. Corticotropin-releasing factor-dopamine interactions in male and female macaque: Beyond the classic VTA. Synapse 2023: 1–20.

40. Biancardi V, Saini J, Pageni A, Prashaad MH, Funk GD, Pagliardini S. Mapping of the excitatory, inhibitory, and modulatory afferent projections to the anatomically defined active expiratory oscillator in adult male rats. The Journal of comparative neurology 2021; 529(4): 853–884.

41. Cho YT, Fudge JL. Heterogeneous dopamine populations project to specific subregions of the primate amygdala. Neuroscience 2010; 165(): 1501–1518.

42. Mabry SJ, McCollum LA, Farmer CB, Bloom ES, Roberts RC. Evidence for altered excitatory and inhibitory tone in the post-mortem substantia nigra in schizophrenia. World J Biol Psychiatry 2020; 21(5): 339–356.

43. Selemon LD, Begovic A. Reduced Midbrain Dopamine Neuron Number in the Adult Non-human Primate Brain after Fetal Radiation Exposure. Neuroscience 2020; 442: 193–201.

44. Totty MS, Cervera Juanes R, Bach SV, Ben Ameur L, Valentine MR, Simons E et al. Transcriptomic diversity of amygdalar subdivisions across humans and nonhuman primates. Sci Adv 2025; 11(38): eadw1029.

45. Roy B, Jacobson A. The intimate relationships of mRNA decay and translation. Trends Genet 2013; 29(12): 691–699.

46. Conrad WS, Oriol L, Kollman GJ, Faget L, Hnasko TS. Proportion and distribution of neurotransmitter-defined cell types in the ventral tegmental area and substantia nigra pars compacta. Addiction Neuroscience 2024; 13.

47. Yamaguchi T, Qi J, Wang HL, Zhang S, Morales M. Glutamatergic and dopaminergic neurons in the mouse ventral tegmental area. The European journal of neuroscience 2015; 41(6): 760–772.

48. Yamaguchi T, Sheen W, Morales M. Glutamatergic neurons are present in the rat ventral tegmental area. The European journal of neuroscience 2007; 25(1): 106–118.

49. Yamaguchi T, Wang HL, Morales M. Glutamate neurons in the substantia nigra compacta and retrorubral field. The European journal of neuroscience 2013; 38(11): 3602–3610.

50. Trudeau LE, Hnasko TS, Wallen-Mackenzie A, Morales M, Rayport S, Sulzer D. The multilingual nature of dopamine neurons. Progress in brain research 2014; 211: 141–164.

51. Ma S, Zhong H, Liu X, Wang L. Spatial Distribution of Neurons Expressing Single, Double, and Triple Molecular Characteristics of Glutamatergic, Dopaminergic, or GABAergic Neurons in the Mouse Ventral Tegmental Area. J Mol Neurosci 2023; 73(6): 345–362.

52. Morales M, Root DH. Glutamate Neurons within the Midbrain Dopamine Regions. Neuroscience 2014; 282: 60–68.

53. Li X, Qi J, Yamaguchi T, Wang HL, Morales M. Heterogeneous composition of dopamine neurons of the rat A10 region: molecular evidence for diverse signaling properties. Brain structure & function 2013; 218(5): 1159–1176.

54. Descarries L, Berube-Carriere N, Riad M, Bo GD, Mendez JA, Trudeau LE. Glutamate in dopamine neurons: synaptic versus diffuse transmission. Brain research reviews 2008; 58(2): 290–302.

55. Wang F, Flanagan J, Su N, Wang LC, Bui S, Nielson A et al. RNAscope: a novel in situ RNA analysis platform for formalin-fixed, paraffin-embedded tissues. The Journal of molecular diagnostics : JMD 2012; 14(1): 22–29.

56. Root DH, Barker DJ, Estrin DJ, Miranda-Barrientos JA, Liu B, Zhang S et al. Distinct Signaling by Ventral Tegmental Area Glutamate, GABA, and Combinatorial Glutamate-GABA Neurons in Motivated Behavior. Cell reports 2020; 32(9): 108094.

57. Barbano MF, Wang H, Zhang S, Shevelkin AV, Yu KJ, Richie CT et al. VTA monosynaptic connections by local glutamate and GABA neurons and their distinct roles in behavior. Nature communications 2025; 16(1): 8500.

58. Wang HL, Qi J, Zhang S, Wang H, Morales M. Rewarding Effects of Optical Stimulation of Ventral Tegmental Area Glutamatergic Neurons. The Journal of neuroscience : the official journal of the Society for Neuroscience 2015; 35(48): 15948–15954.

59. Mendez JA, Bourque MJ, Dal Bo G, Bourdeau ML, Danik M, Williams S et al. Developmental and target-dependent regulation of vesicular glutamate transporter expression by dopamine neurons. The Journal of neuroscience : the official journal of the Society for Neuroscience 2008; 28(25): 6309–6318.

60. Berube-Carriere N, Riad M, Dal Bo G, Levesque D, Trudeau LE, Descarries L. The dual dopamine-glutamate phenotype of growing mesencephalic neurons regresses in mature rat brain. The Journal of comparative neurology 2009; 517(6): 873–891.

61. Buck SA, Mabry SJ, Glausier JR, Banks-Tibbs T, Ward C, Kozel J et al. Aging disrupts the coordination between mRNA and protein expression in mouse and human midbrain. Molecular psychiatry 2025; 30(7): 3039–3054.

62. Sulzer D, Joyce MP, Lin L, Geldwert D, Haber SN, Hattori T et al. Dopamine neurons make Glutamatergic synapses in vitro. J Neurosci 1998; 18(12): 4588–4602.

63. Steinkellner T, Zell V, Farino ZJ, Sonders MS, Villeneuve M, Freyberg RJ et al. Role for VGLUT2 in selective vulnerability of midbrain dopamine neurons. J Clin Invest 2018; 128(2): 774–788.

64. Emborg ME, Ma SY, Mufson EJ, Levey AI, Taylor MD, Brown WD et al. Age-related declines in nigral neuronal function correlate with motor impairments in rhesus monkeys. Journal of Comparative Neurology 1998; 401(2): 253–265.

65. Chuhma N, Zhang H, Masson J, Zhuang X, Sulzer D, Hen R et al. Dopamine neurons mediate a fast excitatory signal via their glutamatergic synapses. The Journal of neuroscience : the official journal of the Society for Neuroscience 2004; 24(4): 972–981.

66. Root DH, Mejias-Aponte CA, Zhang S, Wang HL, Hoffman AF, Lupica CR et al. Single rodent mesohabenular axons release glutamate and GABA. Nature neuroscience 2014; 17(11): 1543–1551.

67. Stuber GD, Hnasko TS, Britt JP, Edwards RH, Bonci A. Dopaminergic terminals in the nucleus accumbens but not the dorsal striatum corelease glutamate. The Journal of neuroscience : the official journal of the Society for Neuroscience 2010; 30(24): 8229–8233.

68. Tecuapetla F, Patel JC, Xenias H, English D, Tadros I, Shah F et al. Glutamatergic signaling by mesolimbic dopamine neurons in the nucleus accumbens. The Journal of neuroscience : the official journal of the Society for Neuroscience 2010; 30(20): 7105–7110.

69. Zigmond MJ. Chemical transmission in the brain: homeostatic regulation ands its functional implications. In: Bloom F (ed). Neuroscience: From the Molecular to the Cognitive. Elsevier Science B.V. 1994, pp 115–122.

70. Tank AW, Xu L, Chen X, Radcliffe P, Sterling CR. Post-transcriptional regulation of tyrosine hydroxylase expression in adrenal medulla and brain. Annals of the New York Academy of Sciences 2008; 1148: 238–248.

71. Eiden LE, Weihe E. VMAT2: a dynamic regulator of brain monoaminergic neuronal function interacting with drugs of abuse. Annals of the New York Academy of Sciences 2011; 1216: 86–98.

72. Srinivasan S, Limani F, Hanzlova M, La Batide-Alanore S, Klotz S, Hnasko TS et al. Evidence for low affinity of GABA at the vesicular monoamine transporter VMAT2 - Implications for transmitter co-release from dopamine neurons. Neuropharmacology 2025; 270: 110367.

73. Kim JI, Ganesan S, Luo SX, Wu YW, Park E, Huang EJ et al. Aldehyde dehydrogenase 1a1 mediates a GABA synthesis pathway in midbrain dopaminergic neurons. Science 2015; 350(6256): 102–106.

74. Tritsch NX, Oh WJ, Gu C, Sabatini BL. Midbrain dopamine neurons sustain inhibitory transmission using plasma membrane uptake of GABA, not synthesis. eLife 2014; 3: e01936.

75. Stamatakis AM, Jennings JH, Ung RL, Blair GA, Weinberg RJ, Neve RL et al. A unique population of ventral tegmental area neurons inhibits the lateral habenula to promote reward. Neuron 2013; 80(4): 1039–1053.

76. Tiklova K, Bjorklund AK, Lahti L, Fiorenzano A, Nolbrant S, Gillberg L et al. Single-cell RNA sequencing reveals midbrain dopamine neuron diversity emerging during mouse brain development. Nature communications 2019; 10(1): 581.

77. Eskenazi D, Malave L, Mingote S, Yetnikoff L, Ztaou S, Velicu V et al. Dopamine Neurons That Cotransmit Glutamate, From Synapses to Circuits to Behavior. Front Neural Circuits 2021; 15: 665386.

78. Hnasko TS, Edwards RH. Neurotransmitter corelease: mechanism and physiological role. Annu Rev Physiol 2012; 74: 225–243.

79. Poulin JF, Gaertner Z, Moreno-Ramos OA, Awatramani R. Classification of Midbrain Dopamine Neurons Using Single-Cell Gene Expression Profiling Approaches. Trends in neurosciences 2020; 43(3): 155–169.

80. Farassat N, Costa KM, Stojanovic S, Albert S, Kovacheva L, Shin J et al. In vivo functional diversity of midbrain dopamine neurons within identified axonal projections. eLife 2019; 8.

81. Mingote S, Amsellem A, Kempf A, Rayport S, Chuhma N. Dopamine-glutamate neuron projections to the nucleus accumbens medial shell and behavioral switching. Neurochem Int 2019; 129: 104482.

82. Poulin JF, Caronia G, Hofer C, Cui Q, Helm B, Ramakrishnan C et al. Mapping projections of molecularly defined dopamine neuron subtypes using intersectional genetic approaches. Nature neuroscience 2018; 21(9): 1260–1271.

83. Yamaguchi T, Wang HL, Li X, Ng TH, Morales M. Mesocorticolimbic glutamatergic pathway. The Journal of neuroscience : the official journal of the Society for Neuroscience 2011; 31(23): 8476–8490.

84. Gorelova N, Mulholland PJ, Chandler LJ, Seamans JK. The glutamatergic component of the mesocortical pathway emanating from different subregions of the ventral midbrain. Cereb Cortex 2012; 22(2): 327–336.

85. Taylor SR, Badurek S, Dileone RJ, Nashmi R, Minichiello L, Picciotto MR. GABAergic and glutamatergic efferents of the mouse ventral tegmental area. The Journal of comparative neurology 2014; 522(14): 3308–3334.

86. Mingote S, Chuhma N, Kusnoor SV, Field B, Deutch AY, Rayport S. Functional Connectome Analysis of Dopamine Neuron Glutamatergic Connections in Forebrain Regions. The Journal of neuroscience : the official journal of the Society for Neuroscience 2015; 35(49): 16259–16271.

87. Haber SN, Fudge JL, McFarland N. Striatonigrostriatal pathways in primates form an ascending spiral from the shell to the dorsolateral striatum. J Neurosci 2000; 20: 2369–2382.

88. Hedreen JC, DeLong MR. Organization of striatopallidal, striatonigal, and nigrostriatal projections in the Macaque. J Comp Neurol 1991; 304: 569–595.

89. Raghanti MA, Stimpson CD, Marcinkiewicz JL, Erwin JM, Hof PR, Sherwood CC. Cortical dopaminergic innervation among humans, chimpanzees, and macaque monkeys: a comparative study. Neuroscience 2008; 155(1): 203–220.

90. Berger B, Gaspar P, Verney C. Dopaminergic innervation of the cerebral cortex: unexpected differences between rodents and primates [published erratum appears in Trends Neurosci 1991 Mar;14(3):119]. Trends in neurosciences 1991; 14(1): 21–27.

91. Gaspar P, Stepneiwska I, Kaas JH. Topography and collateralization of the dopaminergic projections to motor and lateral prefrontal cortex in owl monkeys. J Comp Neurol 1992; 325: 1–21.

92. Williams SM, Goldman-Rakic PS. Widespread origin of the primate mesofrontal dopamine system. Cerebral Cortex 1998; 8(4): 321–345.

93. Matsumoto M, Hikosaka O. Two types of dopamine neuron distinctly convey positive and negative motivational signals. Nature 2009; 459(7248): 837–841.

94. La Manno G, Gyllborg D, Codeluppi S, Nishimura K, Salto C, Zeisel A et al. Molecular Diversity of Midbrain Development in Mouse, Human, and Stem Cells. Cell 2016; 167(2): 566–580 e519.

95. Fortin GM, Bourque MJ, Mendez JA, Leo D, Nordenankar K, Birgner C et al. Glutamate corelease promotes growth and survival of midbrain dopamine neurons. The Journal of neuroscience : the official journal of the Society for Neuroscience 2012; 32(48): 17477–17491.

96. Schmitz Y, Luccarelli J, Kim M, Wang M, Sulzer D. Glutamate controls growth rate and branching of dopaminergic axons. The Journal of neuroscience : the official journal of the Society for Neuroscience 2009; 29(38): 11973–11981.

97. Avramescu RG, Hernandez G, Flores C. Rewiring the future: drugs abused in adolescence may predispose to mental illness in adult life by altering dopamine axon growth. J Neural Transm (Vienna) 2024; 131(5): 461–467.

98. Rosenberg DR, Lewis DA. Postnatal maturation of the dopaminergic innervation of monkey prefrontal and motor cortices: A tyrosine hydroxylase immunohistochemical analysis. J Comp Neurol 1995; 358: 383–400.

99. Reynolds LM, Flores C. Mesocorticolimbic Dopamine Pathways Across Adolescence: Diversity in Development. Front Neural Circuits 2021; 15: 735625.

100. Islam KUS, Meli N, Blaess S. The Development of the Mesoprefrontal Dopaminergic System in Health and Disease. Front Neural Circuits 2021; 15: 746582.

