## Supplementary figures and images for "Multiplexed dopamine neurons predominate in the ventral midbrain of young macaques"

### Suppl Figure 1

A

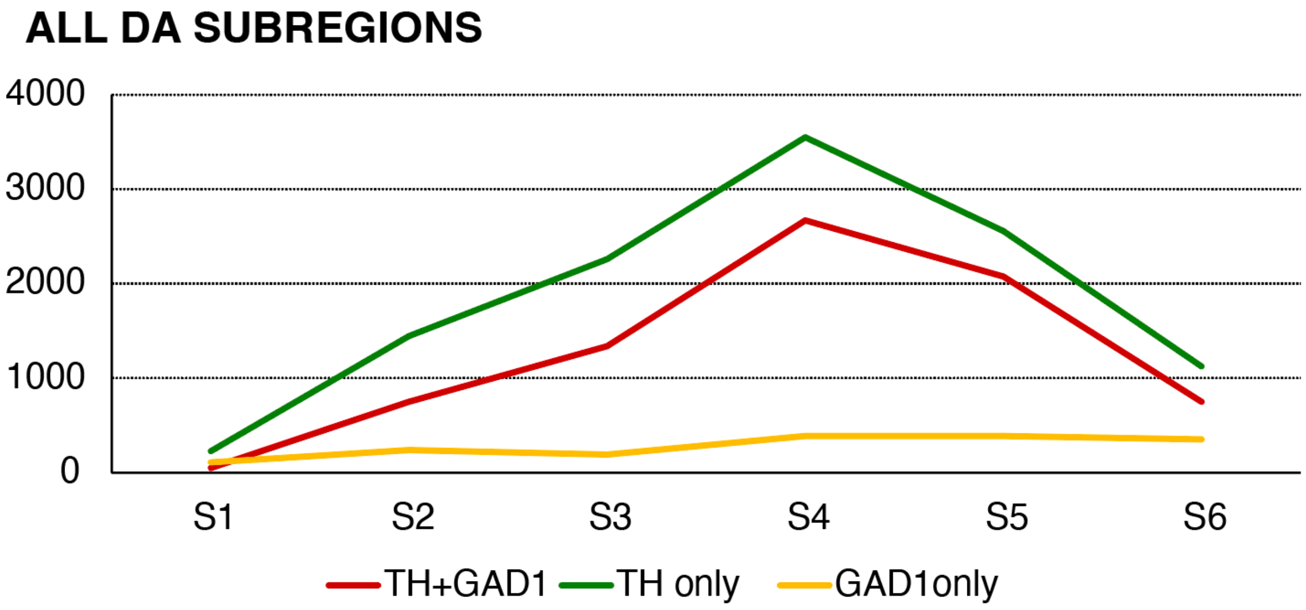


**ALL DA SUBREGIONS**

| | S1 | S2 | S3 | S4 | S5 | S6 |
|---|---|---|---|---|---|---|
| TH+GAD1 | 47 | 748 | 1340 | 2670 | 1683 | 750 |
| TH alone | 228 | 1447 | 2259 | 3551 | 1816 | 1122 |
| GAD1alone | 109 | 239 | 189 | 390 | 306 | 349 |

B

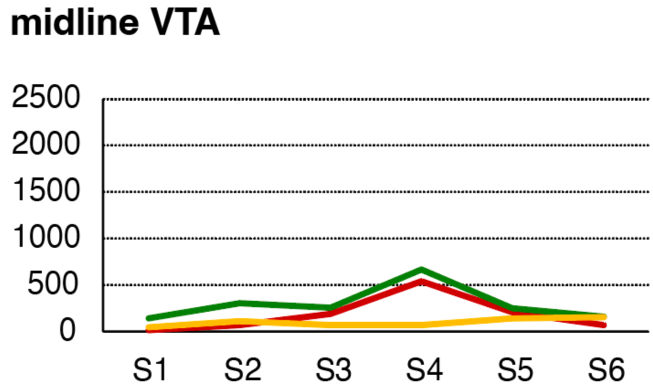


C

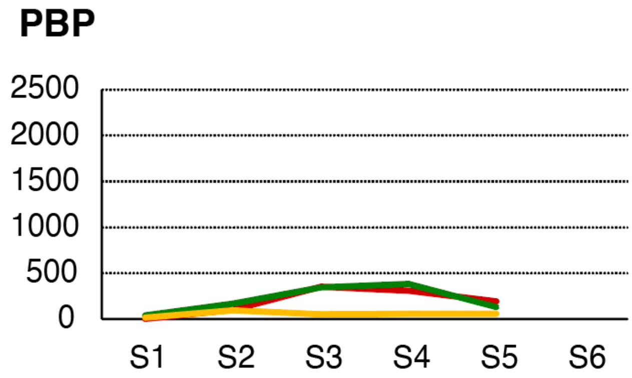


D

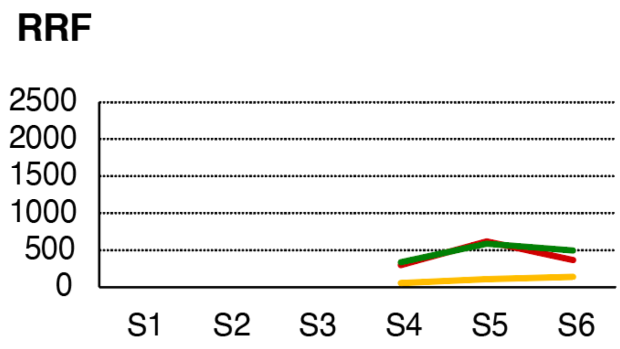


E

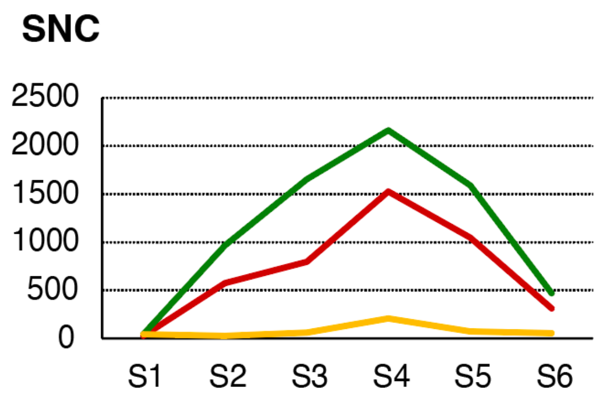
